# StackPDB: predicting DNA-binding proteins based on XGB-RFE feature optimization and stacked ensemble classifier

**DOI:** 10.1101/2020.08.24.264267

**Authors:** Qingmei Zhang, Peishun Liu, Yu Han, Yaqun Zhang, Xue Wang, Bin Yu

## Abstract

DNA binding proteins (DBPs) not only play an important role in all aspects of genetic activities such as DNA replication, recombination, repair, and modification but also are used as key components of antibiotics, steroids, and anticancer drugs in the field of drug discovery. Identifying DBPs becomes one of the most challenging problems in the domain of proteomics research. Considering the high-priced and inefficient of the experimental method, constructing a detailed DBPs prediction model becomes an urgent problem for researchers. In this paper, we propose a stacked ensemble classifier based method for predicting DBPs called StackPDB. Firstly, pseudo amino acid composition (PseAAC), pseudo position-specific scoring matrix (PsePSSM), position-specific scoring matrix-transition probability composition (PSSM-TPC), evolutionary distance transformation (EDT), and residue probing transformation (RPT) are applied to extract protein sequence features. Secondly, extreme gradient boosting-recursive feature elimination (XGB-RFE) is employed to gain an excellent feature subset. Finally, the best features are applied to the stacked ensemble classifier composed of XGBoost, LightGBM, and SVM to construct StackPDB. After applying leave-one-out cross-validation (LOOCV), StackPDB obtains high ACC and MCC on PDB1075, 93.44% and 0.8687, respectively. Besides, the ACC of the independent test datasets PDB186 and PDB180 are 84.41% and 90.00%, respectively. The MCC of the independent test datasets PDB186 and PDB180 are 0.6882 and 0.7997, respectively. The results on the training dataset and the independent test dataset show that StackPDB has a great predictive ability to predict DBPs.

## 1. Introduction

DNA binding proteins (DBPs) are proteins that can bind and interact with DNA and participate in many biological processes [1]. For example, transcription factors participate in the DNA transcription process while nucleases can cut DNA molecules. Besides, histones are related to the packaging of chromatin in the nucleus [2]. DBPs are essential components of anticancer drugs, antibiotics, and steroids in the research of anticancer drugs and the treatment of genetic diseases. Meanwhile, DBPs have an irreplaceable role in the biophysical, biochemical, and biological research of DNA [3]. Early identification of DBPs generally used experimental methods such as filter combining analysis [4], genetic analysis [5], chromatin immunoprecipitation [6], and X-ray crystallography [7]. With the deep research of high-throughput sequencing technology, protein sequences continue to emerge. However, traditional biological experiment methods are time-consuming and expensive. Identifying DBPs based on experimental methods that are far from meeting the research needs [8]. Therefore, computational methods are used as powerful tools to predict DBPs.

Researchers have developed numerous calculation methods to identify DBPs. The important step of predicting DBPs is to extract features from protein sequences. Feature extraction methods can dig four types of protein sequence information which are sequence information, physicochemical properties, structural information, and evolutionary information. Rahman et al. [9] used amino acid composition (AAC), dipeptides composition (DC), tripeptides composition (TC), n-gapped-dipeptides (nGDip), and position-specific n-grams (PSN) to obtain protein sequence information. Zhang et al. [10] used 14 kinds of physicochemical property, protein secondary structural information, and evolutionary information to predict DBPs. Chowdhury et al. [11] used PSI-BLAST to obtain the PSSM, which indicated the evolutionary information. SPIDER2 was used to extract the secondary structural information of the protein sequences. Nanni et al. [12] used AAC and quasi residue couple (QRC) to extract protein sequence information. Meanwhile, physicochemical properties were extracted by the autocovariance approach (AC). In addition, pseudo-position specific scoring matrix (PsePSSM), N-gram features (NGR) and texture descriptors (TD) extracted evolutionary information. Sang et al. [13] obtained the HMM matrix according to the hidden Markov model (HMM) for each sequence. AAC, autocovariance transformation (ACT), and cross-covariance transformation (CCT) were used to convert the HMM matrix into feature vectors of the same length. Then DBPs prediction was performed after fusing multiple features.

Although the fusion of multiple features can fully represent the information contained in the protein sequence, it may also bring redundancy and noise that will reduce the efficiency of the model. Therefore, choosing an appropriate dimension reduction method is also an important step in the process of DBPs identification. Hu et al. [14] fused four feature extraction methods of AAC, pseudo predicted relative solvent accessibility (PsePRSA), PsePSSM, and pseudo predicted probabilities of DNA-binding sites (PsePPDBS). Support vector machine recursive feature elimination and correlation bias reduction (SVM-RFE+CBR) [15] was used to convert the nonlinear learning issue in the original feature space to a linear learning issue in the high dimension feature space. The optimal feature subset containing 131-dimension vectors was obtained by SVM-RFE+CBR. Zhou et al. [16] used dipeptide deviation from the expected mean (DDE), normalized Moreau-broto autocorrelation (NMBAC), PSSM-distance-bigram transformation (PSSM-DBT), and PSSM-discrete wavelet transformation (PSSM-DWT) to extract features. After fusing the obtained features, SVM-RFE+CBR was used for dimension reduction to obtain a feature subspace containing 424-dimension vectors. Ali et al. [17] performed feature extraction based on PSSM, PSSM-DWT, and split amino acid composition (SAAC). Then they used maximum relevance and minimum redundancy (mRMR) to decrease the number of fused features. mRMR sorted each feature in the feature space according to the maximum relevance and minimum redundancy with the target class, and finally obtained the optimal subset containing 264-dimension features. Ji et al. [18] adopted AAC, DC, chaos game representation (CGR), fractal dimension (FD), composition transition and distribution (CTD), Moreau-Broto (MB), PseAAC, sequence order (SO) and PSSM to extract features of the training dataset. Multi-class MSVM-RFE was used for dimension reduction. MSVM-RFE converted the multi-objective optimization issue to a single-objective optimization issue. The redundant features are gradually removed according to the sorting criteria, and the optimal subset containing 100-dimension features is obtained.

In addition to choosing appropriate feature extraction and feature selection algorithms, another key factor for the success of DBPs prediction is the choice of classification algorithms. Appropriate classification algorithms can efficiently shorten the running time and learn the relationship between tags and categories. Some machine learning methods are commonly used, such as K Nearest Neighbor (KNN) [19], Neural Network [20], Navïe Bayes [21], Hidden Markov Model [22], Gradient Boosting Decision Tree (GBDT) [23], Support Vector Machine (SVM) [24] and (RF) [25] and etc. Ali et al. [26] proposed the DP-BINDER model. According to the feature selection method SVM-RFE+CBR, 84-dimension features were input into RF and SVM for prediction. Based on the LOOCV, the prediction accuracy of the training dataset PDB1075 reached 92.46% and 91.72%, respectively. Kumar et al. [27] used amino acid and dipeptide composition, PSSM-400, four-part amino acid composition for feature extraction. Additionally, SVM was used for prediction. The ACC of the model reached 74.22%. Wei et al. [28] proposed the Local-DPP model, which used Local PsePSSM to get the local protection information. Taking the obtained 120-dimension feature vectors as the input of RF, the ACC of the Local-DPP model over the LOOCV reached 79.2%. Chauhan et al. [29] added 0 vectors to the PSSM to generate a fixed-length padded matrix (pPSSM) and then used deep convolutional neural networks (CNNs) to predict DBPs. Liu et al. [30] proposed the MFSBinder method, which used Local-DPP, 188D, PSSM-DWT, and AC-struct to extract evolutionary information, sequence information, physicochemical properties, and structural information, respectively. Finally, a stacked ensemble classifier was used to predict DBPs. Xu et al. [31] extracted physicochemical property, amino acid composition and distribution information. Then the features were used to predict DBPs based on unbalanced-AdaBoost. Liu et al. [32] proposed the iDNA-KACC model which combined contour-based protein expression, self-crossing covariance transformation, and Kmer composition features. The features were fed to an ensemble classifier composed of 4 SVMs for prediction. The ACC of the iDNA-KACC model was 75.16% based on LOOCV.

Although the existing methods can effectively predict DBPs, the running speed and accuracy of the methods need to be improved. First, the influence of protein sequence features on DBPs prediction has not been fully elucidated. It still has to be improved in DBPs prediction by extracting features based on protein sequences. Second, feature fusion brings redundancy and noise. Choosing a suitable dimension reduction method can reduce the feature dimension while retaining effective information. Finally, since the number of protein sequences continuously increase, choosing an effective classifier is also a major challenge for researchers.

Hence, we proposed a new DBPs prediction model, called StackPDB. Firstly, the training dataset PDB1075 was encoded into EDT, RPT, PseAAC, PsePSSM, and PSSM-TPC. Compared with the individual feature, the fusion feature can obtain more comprehensive protein information. Secondly, we applied XGB_RFE to the DBPs prediction field for the first time. XGB_RFE can speed up the process of the StackPDB model and choose the best features while deleting irrelevant features and reducing the feature dimension. Finally, the stacked ensemble classifier was used as the final classifier. In the first stage, two XGBoost and two LightGBM were used for the first time. Then the output probability of the base-classifier was input into the meta-classifier SVM for DBPs prediction. The ACC of StackPDB on the training dataset PDB1075 reached 93.44% over the LOOCV test. Using the independent test datasets PDB186 and PDB180 to test the generalization ability of the StackPDB model, StackPDB obtained an ACC value of 84.40% and 90.00%, respectively. Compared with other competitive methods, StackPDB has higher stability and can significantly improve the recognition ability of DBPs.

## 2. Materials and methods

### 2.1. Datasets

Choosing the appropriate data set is a key step to build a model. In this article, we chose the dataset PDB1075 as the training dataset. Xu et al. [33] established the training dataset PDB1075 which contains 525 DBPs and 550 non-DBPs. The dataset construction process met the following criteria: (1) Searching from the updated protein database (PDB) to acquire DBPs sequences; (2) Protein sequences that less than 50 in length or contained the character “X” were removed; and (3) Sequences with sequence similarity greater than 25% in the same dataset were removed by the software PISCES. During the experiment in this article, we found 8 abnormal sequences in the training dataset: (1) 1AOII, (2) 4FCYC, (3) 4JJNJ, (4) 4JJNI, (5) 3THWD, (6) 4GNXL, (7) 4GNXZ, (8) 2RAUA, where the first four were DNA sequences, and the PSSM matrix of the last four sequences were not available in the PSI-BLAST [34] program. After deleting abnormal sequences, the training dataset consists of 518 DBPs and 549 non-DBPs were used in this article.

To test our model, we chose PDB186 and PDB180 as independent test datasets. The independent test dataset PDB186 was collected by Lou et al. [35] which contains 93 DBPs and 93 non-DBPs. The independent test dataset PDB180 was proposed by Xu et al. [36] which contains 81 DBPs and 99 non-DBPs. The two independent test sets used the same processing method in the construction process. During the construction of two independent test sets, length of protein sequences less than 60 or the character “X” were removed. BLASTCLUST software was used to remove sequences with a sequence similarity greater than 25% in the same dataset.

### 2.2. Feature extraction

#### 2.2.1. Pseudo amino acid composition (PseAAC)

Chou [37] proposed PseAAC, which extracted protein sequence and physicochemical information. PseAAC has been applied in many fields, e.g., the subcellular location of apoptosis proteins [38], protein structural prediction [39], protein post-translational modification site prediction [40], protein submitochondrial localization prediction [41] and etc.

The feature vector is obtained by PseAAC as follows:

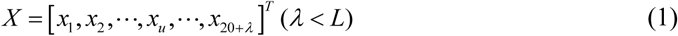

The calculation method *x*_*u*_ is shown in formula (2)

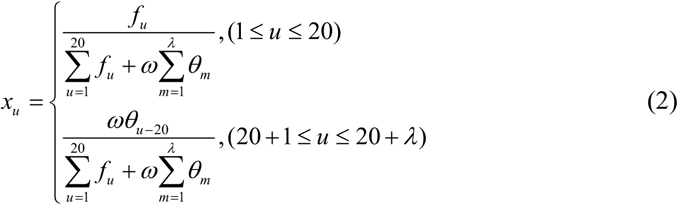

where *L* represents the length of the protein sequence while *f*_*u*_ is the frequency of the *u*- th amino acid in the protein sequence *S*. *θ*_*m*_ is the *m* -layer sequence correlation factor. *ω* is the weighting factor where *ω* = 0.05. PseAAC extracts 20 + *λ* -dimension feature vectors. The first 20-dimension vectors represent amino acid sequence information, and the latter *λ* -dimension represents amino acid sequence order information and physicochemical properties.

#### 2.2.2. Position-specific scoring matrix (PSSM)

Evolutionary information is vital information in protein function annotation. It has been widely used in many fields, such as protein-protein interaction prediction [42], RNA-protein interaction prediction [43], DNA binding proteins prediction [44] and etc. In this paper, PsePSSM, PSSM-TPC, EDT, and RPT are used to extract evolutionary information. The four feature extraction methods are based on the PSSM, so PSSM is initially introduced. Jones et al. [45] firstly proposed PSSM, using the PSI-BLAST [34] program to perform three iterative searches in the Swiss-Prot database, and the *E* value threshold was set as 0.001. By performing multiple sequences comparisons on protein sequences, a *L* × 20 PSSM is generated, as shown in formula (3).

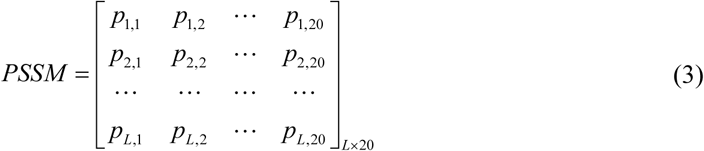

where *p*_*i, j*_ represents the score of the *i* -th amino acid mutates into the *j* -th standard amino acid during the evolution process. *L* represents the length of the protein sequence. To eliminate the dimensional error, the PSSM is standardized according to formula (4):

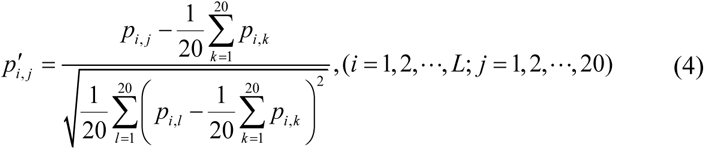

Where 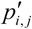 represents the PSSM element after standardization. PSSM is changed to a vector with equal length by formula (5-6).

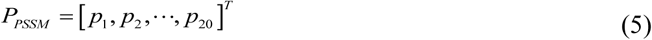

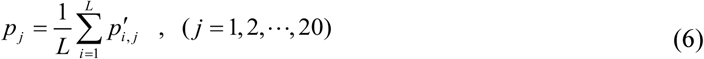

where *P*_*PSSM*_ represents a feature vector of length 20 and *p*_*j*_ represents a vector element.

#### 2.2.3. Pseudo-position specific scoring matrix (PsePSSM)

Although *P*_*PSSM*_ contains evolutionary information, it ignores the sequence order information. At present, PsePSSM [46] has been applied to human protein subcellular localization identification [47], protein submitochondrial localization [48], drug-target interaction prediction [49], membrane protein recognition [50] and etc. PsePSSM is shown in equation (7).

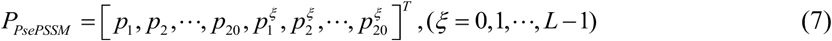

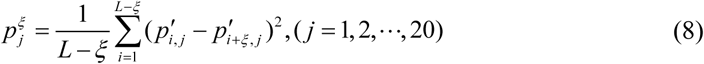

where 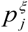 represents the correlation of the PSSM score between two amino acids separated by *ξ*. The accuracy of prediction is changed by adjusting *ξ*. We took *ξ* from 1 to 9 with 1 as the interval and determined the optimal *ξ* value of 2. According to PsePSSM, 20+20×*ξ* = 60 -dimension feature vectors can be obtained for each protein sequence.

#### 2.2.4. Position-specific scoring matrix-transition probability composition (PSSM-TPC)

To reduce the loss of sequence information in the evolution process, transition probability composition (TPC) is applied to PSSM. The procedure given in [51] is used to calculated TPC by the transition probability matrix (TPM). The PSSM-TPC vector can be expressed by formula (9):

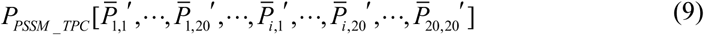

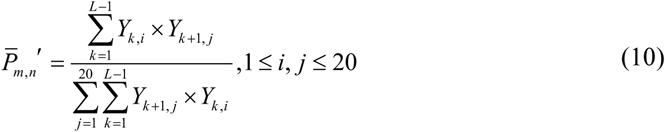

where 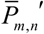 represents the transition probability from the *m* -th amino acid to the *n* -th amino acid. *Y*_*i, j*_ which satisfies 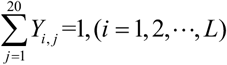 represents the relative probability of the *j* -th amino acid appearing at the *i* -th position.

#### 2.2.5. Evolutionary distance transformation (EDT)

EDT was proposed by Zhang et al. [52] which calculated the non-co-occurrence probability of two amino acids. The amino acids are separated by *d* (*d* = 1, 2,…, *L*_min_ −1). EDT can be calculated by the formula (11):

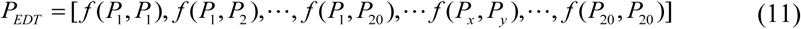

The non-co-occurrence probability *f* (*P*_*x*_, *P*_*y*_) of two amino acids separated by *d* can be calculated by the formula (12):

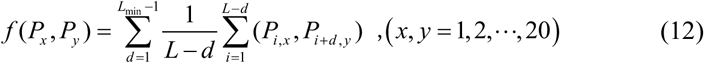

where *L*_min_ represents the minimum sequence length and *P*_*x*_, *P*_*y*_ represents 20 different standard amino acids. *P*_*i, x*_ and *P*_*i* + *d, y*_ are both elements in PSSM. Hence, EDT extracts 400-dimension features representing non-collinear probability information.

#### 2.2.6. Residue probing transformation (RPT)

RPT was proposed by Jeong et al. [53], grouping the evolution scores in the PSSM to emphasize domains with similar conservation. The rows of the same amino acid in the PSSM are divided into one group. Thus, a total of 20 groups are obtained. For each group, the sum of the elements in each column is calculated. In this way, each protein sequence can get an 20 × 20 RPT matrix, as shown in equation (13):

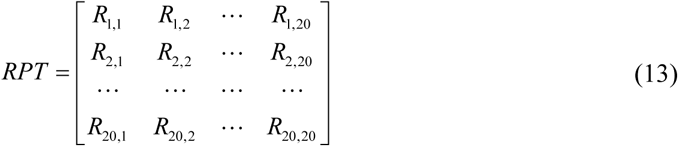

A 400-dimension row vector is obtained by expanding the RPT matrix, as shown in formula (14):

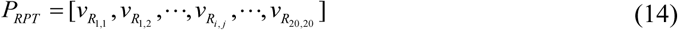

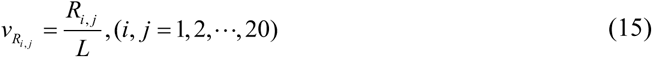

where *R*_*i, j*_ represents the RPT element. *L* is the sequence length, and *P*_*RPT*_ represents the 400-dimension feature vector obtained by RPT.

### 2.3. Extreme gradient boosting-recursive feature elimination (XGB-RFE)

The XGBoost algorithm was proposed by Yu et al. [54], which sorted the input features according to their importance. First, the algorithm uses XGBoost to obtain significance mark of every feature, and assign weights to the features. Then, the weighted sum of the scores of each feature in all boost trees is used to obtain the final importance score. Then the features are sorted according to the final score. In this paper, XGBoost and recursive feature elimination algorithm (RFE) [55] are combined for the first time in the field of DBPs prediction.

Given a set *D* = {(*x*_*i*,1_, *y*_*i*_), (*x*_*i*,2_, *y*_*i*_),, (*x*_*i,m*_, *y*_*i*_)}, the element (*x*_*i,m*_, *y*_*i*_) = (*x*_*i*,1_, *x*_*i*,2_,, *x*_*i,m*_) indicates that the label of *m*-th feature vector is *y*_*i*_.

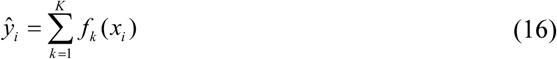

where *f*_*k*_ (*x*_*i*_) represents the importance score of *i* -th feature vector on *k* -th tree.

Then the objective function can be expressed as formula (17):

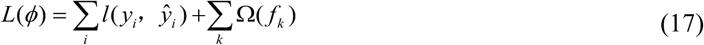

where *l*(*ŷ*_*i*_, *y*_*i*_) represents the loss between the true value and the predicted value. 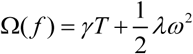 controls the complexity of the model.

Assuming that each iteration can generate a tree, the objective function becomes as follows.

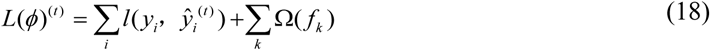

where 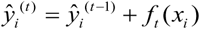 represents the predicted value of *t* -th iteration. Supposing the *k* − 1 -th tree is known while generating the *k* -th tree.

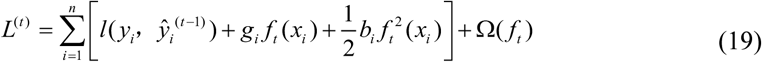

where *L*^(*t*)^ is the objective function. 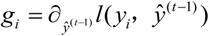 and 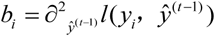 represent the first-order and second-order statistics of the loss function, respectively.

After getting the importance ranking of features, RFE is used to delete the least important features from the current feature space. The process repeats N times until the required number of features is obtained.

### 2.4. Stacked ensemble classifier

The stacked ensemble classifier is an integrated method proposed by Wolpert et al. [56]. The prediction results of multiple ordinary learners are used as new features for retraining. By doing this, the stacked ensemble classifier can achieve the purpose of minimizing the error rate of the prediction model. At present, this method has been applied to predict ncRNA-protein interactions [57], Bacterial Type IV Secreted Effectors [58], anticancer drug response [59], MicroRNA automatic classification [60] and etc. In this paper, a stacked ensemble classifier which including two stages of learning is used to predict DBPs. In the first stage, the features are input into the base-classifier to output the binding probability and non-binding probability of DBPs. In order to enrich the features that are input into the meta-classifier, we chose base-classifier from 9 classifiers, e.g., k-nearest neighbor (KNN) [61], support vector machines (SVM) [62], random forest (RF) [63], gradient boosting decision tree (GBDT) [64], Navïe Bayes classifier (NB) [65], logistic regression (LR) [66], light gradient boosting machine (LightGBM) [67], extreme gradient boosting (XGBoost) [54], and adaptive boosting (AdaBoost) [68]. Finally, XGBoost and LightGBM are selected as the best combination of base-classifier. Then the output results of the first stage input into the meta-classifier. To make full use of the features from the first stage, we chose the best meta-classifier among 9 classifiers, e.g., NB, XGBoost, AdaBoost, LightGBM, KNN, RF, GBDT, LR, and SVM. The prediction results show that the StackPDB model constructed by the meta-classifier SVM and the base-classifier XGBoost and LightGBM is the best. Finally, two XGBoost and two LightGBM are used as the base-classifier, and SVM is our meta-classifier. Algorithm 1 represents the pseudo code of the stacked ensemble classifier.

#### Algorithm 1

Stacked ensemble classifier

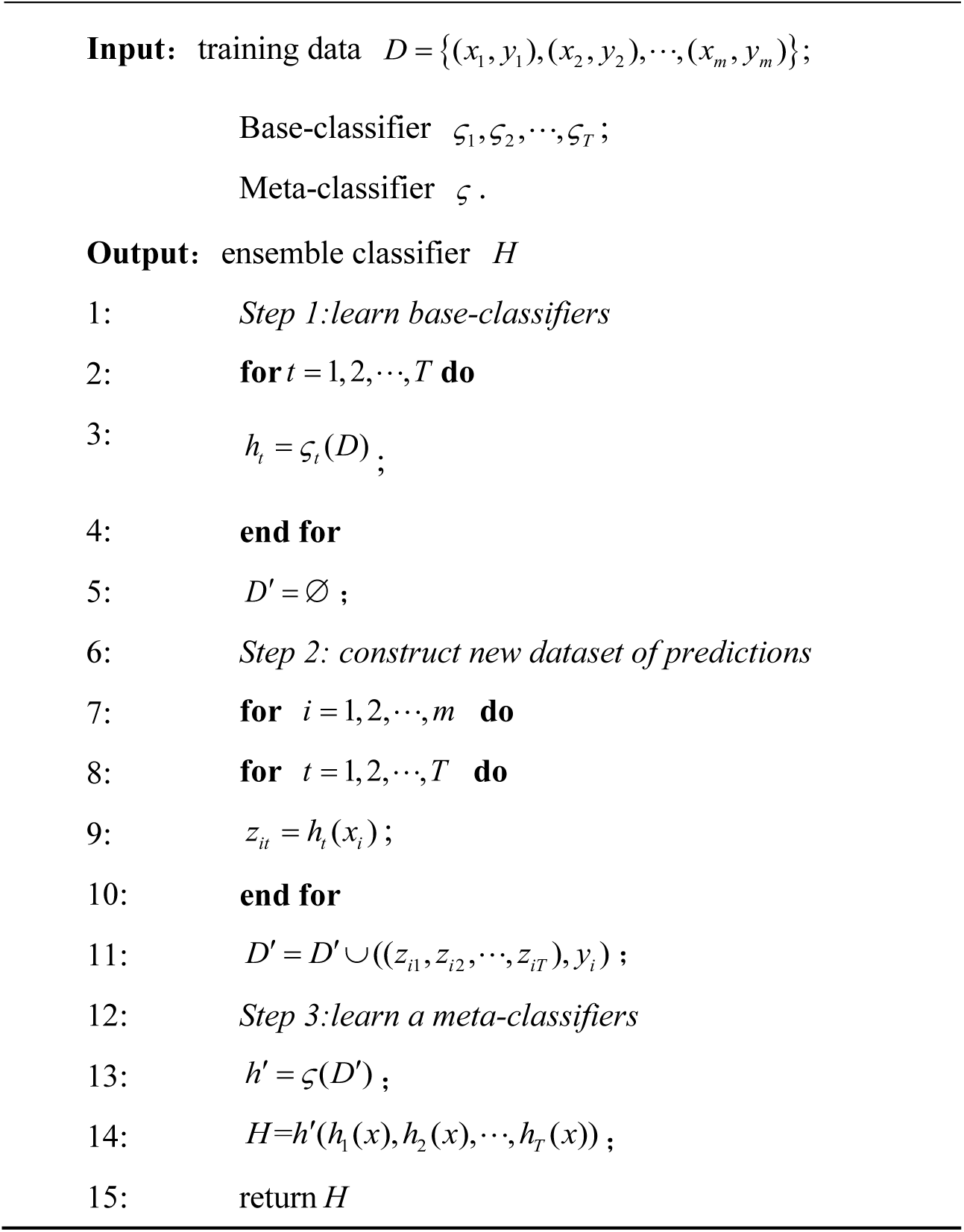

### 2.5. Model construction and evaluation

In this study, we propose a novel model for predicting DBPs, called StackPDB, and the flowchart is shown in Fig. 1. All experiments are performed on Windows Server 2012r 2 Intel (R) Xeon (TM) CPU E5-2650@2.30GHz 2.30GHz, 32.0GB memory, MATLAB2014a, and Python 3.6 programming. The specific algorithm flow is as follows:

**Fig. 1.**
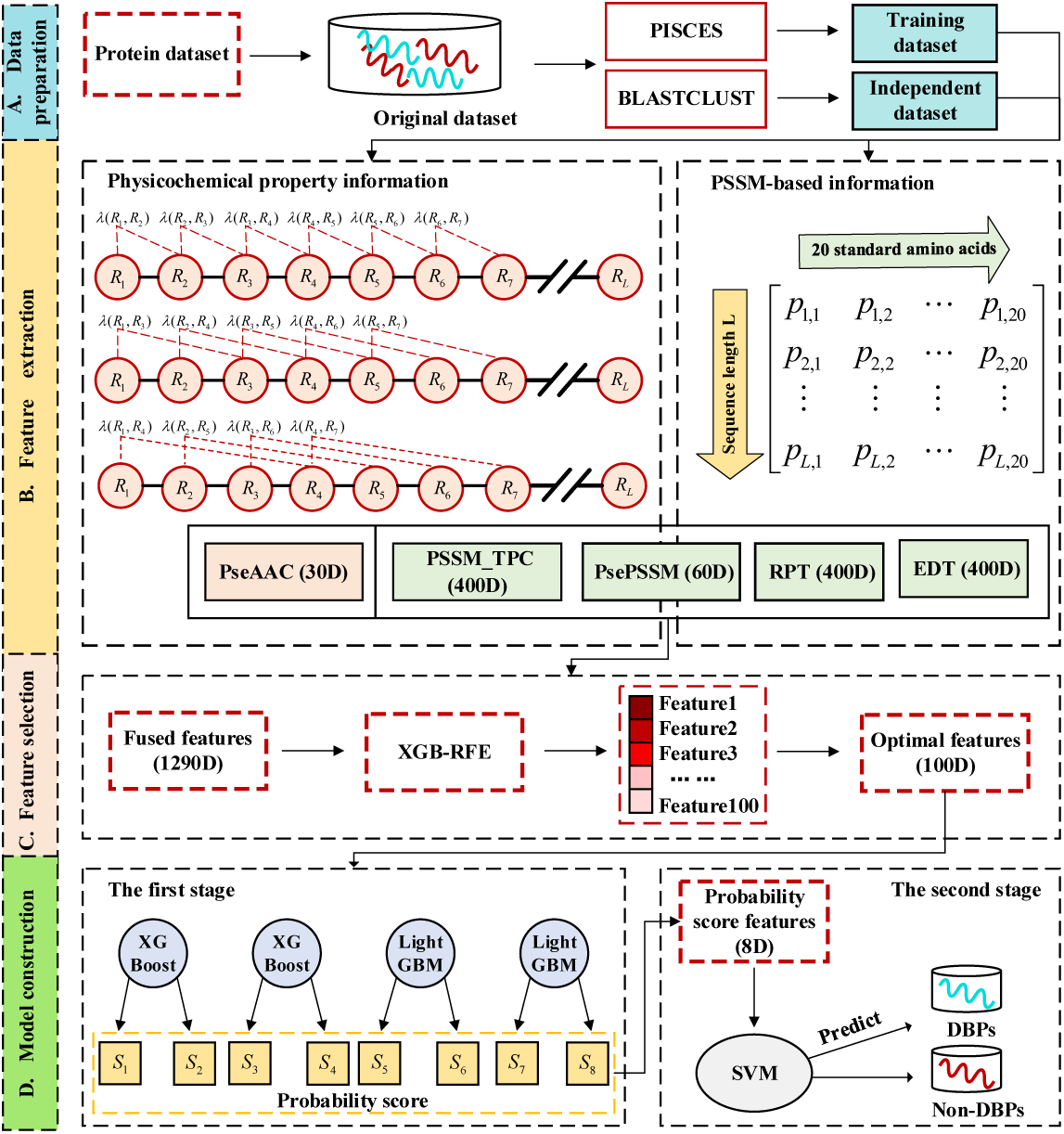
Flow chart of StackPDB. StackPDB firstly collects datasets (A) and then uses five methods to extract protein features (B). StackPDB reduces the dimension of the fusion features (C). Finally stacked ensemble classifier predicts whether the sequence is DBPs or non-DBPs (D).

1. Data preparation. The training dataset PDB1075 and the independent test datasets PDB186 and PDB180 were obtained from the protein database. The protein sequences and their corresponding DBPs labels were entered into StackPDB.
2. Feature extraction. 400-dimension feature vectors were obtained from EDT, RPT, and PSSM-TPC, respectively. 30-dimension and 60-dimension feature vectors were obtained from PseAAC and PsePSSM, respectively. After fusing the five features, an initial feature space that contained 1290-dimension vectors was obtained.
3. Feature selection. The feature selection algorithm XGB-RFE was used to remove the redundancy and noise of the initial feature space in 2). Then 100-dimension optimal feature vectors were obtained.
4. Model construction. The optimal feature vector was input into the base-classifier XGBoost and LightGBM to output the binding probability and non-binding probability of DBPs. The output probability of the base-classifier was input into the meta-classifier SVM to construct the StackPDB.
5. Model verification and evaluation. The effectiveness of StackPDB was tested on the independent test datasets PDB186 and PDB180.

The LOOCV [69], K-fold cross-validation method, and an independent test method are commonly used methods to evaluate the performance of the model. The LOOCV method is chosen as the validation method. In the verification process, LOOCV selects N-1 samples as the training set and one sample as the test set. LOOCV trains N times on the data set to ensure that each sequence is tested. LOOCV can calculate the accuracy of the prediction model objectively and rigorously and test the generalization ability of the model. It has been widely used in proteomics research [70].

Five evaluation indicators are used to evaluates the quality of the model: Accuracy (ACC), Sensitivity (SN), Matthew’s Correlation Coefficient (MCC), and Specificity (SP) [71].

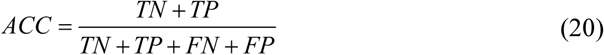

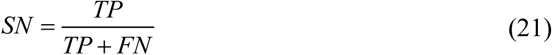

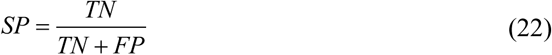

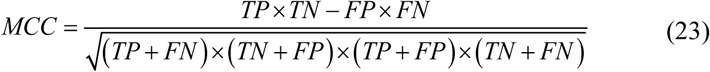

where FN represents the number of DBPs predicted as non-DBPs, FP represents the number of non-DBPs predicted as DBPs, TN represents the number of non-DBPs predicted correctly, and TP represents the number of correct DBPs predicted. Besides, the area under the ROC curve (AUC) and the area under the PR curve (AUPR) are also used as important indicators for evaluating the quality of the model [72, 73].

## 3. Results and discussion

### 3.1. Selection of feature extraction parameters λ and ξ

It is essential to select the excellent parameters when constructing StackPDB model. If the parameter is set too small, the information will be insufficiently extracted. If the parameter is too large, redundant features will be produced. When selecting the best parameter *λ* in PseAAC, the value *λ* is set to 5∼45 with an interval of 5. Similarly, the parameter *ξ* in PsePSSM is set to 1∼10 with an interval of 1. The features with different parameters are used as the input of the stacked ensemble classifier. The prediction results verified by the LOOCV are shown in Supplementary Table S1 and Table S2. The influence of different *λ* of PseAAC and *ξ* of PsePSSM on ACC is shown in Fig. 2.

**Fig. 2.**
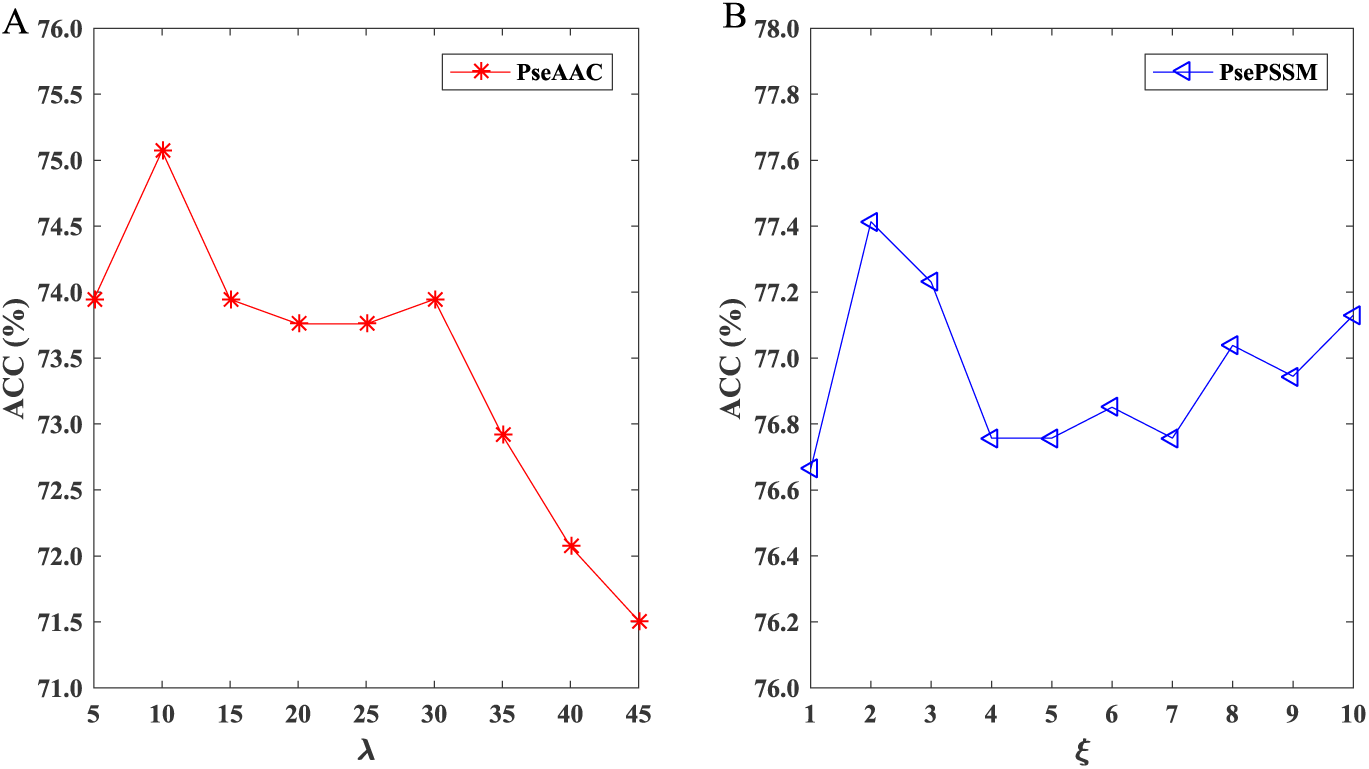
The effect of choosing different *λ* (A) and *ξ* (B) values on the training dataset PDB1075.

In Fig. 2 (A), the performance of PseAAC changes when *λ* gradually increases. The ACC value of PseAAC is the largest when *λ* = 10. As *λ* increasing, the ACC value of StackPDB decreases. As we can see from Fig. 2 (B), the performance of PsePSSM changes when *ξ* increases. The ACC of PsePSSM reaches the maximum value when *ξ* = 2, and then it gradually decreases. When *λ* = 10 the ACC value of PseAAC reaches a maximum of 75.07%. When *ξ* = 2 the ACC of PsePSSM reaches the maximum value of 77.41%, which can fully express protein information. We choose *λ* = 10 as the best parameter of PseAAC so that the PseAAC features can be fully extracted. Finally 20 + *λ* = 30 -dimension feature vectors can be obtained by PseAAC. Similarly, we choose *ξ* = 2 as the best parameter of PsePSSM, so that the PsePSSM features can be fully extracted. Finally 20 + 20×*ξ* = 60 -dimension feature vectors can be obtained by PsePSSM.

### 3.2. Comparison of different feature extraction methods

After determining the best parameters of PseAAC and PsePSSM, EDT, RPT, PseAAC, PsePSSM, and PSSM-TPC are fused to obtain more comprehensive information. To measure the differences between EDT, RPT, PseAAC, PsePSSM, and PSSM-TPC, the 5 individual features and the fusion feature (Fusion) are fed to the stacked ensemble classifier. The results based on the LOOCV are shown in Table 1.

**Table 1.** Performance of 5 feature extraction methods on the training dataset PDB1075.

| Algorithm | ACC (%) | SN (%) | SP (%) | MCC |
| --- | --- | --- | --- | --- |
| EDT | 76.10 | 80.50 | 71.95 | 0.5256 |
| RPT | 76.76 | 83.59 | 70.31 | 0.5425 |
| PseAAC | 75.07 | 74.71 | 73.22 | 0.4791 |
| PsePSSM | 77.41 | 81.85 | 73.22 | 0.5519 |
| PSSM-TPC | 82.48 | 78.98 | 85.79 | 0.6521 |
| Fusion | 89.50 | 87.07 | 91.80 | 0.7903 |

It can be seen from Table 1 that PSSM-TPC performs best among 5 features with an ACC value of 82.48% and an MCC value of 0.6521. The ACC values of EDT, RPT, PseAAC and PsePSSM are 76.10%, 76.76%, 75.07% and 77.41%, respectively, and the MCC values are 0.5256, 0.5425, 0.4791, 0.5519, respectively. For the Fusion features, the value of each evaluation index is improved based on the LOOCV. The MCC and ACC of Fusion are 0.7903 and 89.50%, respectively, which are 13.82% and 7.02% higher than the best single feature PSSM-TPC. Besides, we draw the ROC and PR curves between the single feature extraction method and Fusion as shown in Supplementary Figure S1. The results show that an individual feature can only capture a single aspect of the protein sequence. The Fusion features can obtain more comprehensive information so that it improves the prediction accuracy of DBPs. Nevertheless, multi-information fusion will inevitably bring redundant information.

### 3.3. Comparison of different dimension reduction methods

The dimension reduction method can delete the redundancy while reducing the feature dimension and selecting the optimal feature. After applying fusion of EDT, RPT, PseAAC, PsePSSM, and PSSM-TPC, 1290-dimension feature vectors are obtained. In this paper, 7 feature selection methods are tested on training dataset PDB1075, namely LASSO [47], Elastic net [74], SVM-RFE [26], LinearSVC [75], locally linear embedding (LLE) [76], singular value decomposition (SVD) [77] and XGB_RFE [54]. The parameters are set as follows, (1) The penalty parameter of LASSO is 0.01, thus 197-dimension features are selected; (2) L1_ratio of Elastic net is set to 0.4; (3) SVM-RFE selects the linear kernel function; (4) The penalty of LinearSVC is set to L1; and (5) The optimal features of LLE, SVD and XGB_RFE are set to 100. The final number of features retained by LASSO, Elastic net, SVM-RFE, and LinearSVC are 197, 144, 100, and 386 respectively. The optimal feature subsets obtained by different dimension reduction methods are classified by stacked ensemble classifier. The prediction results are shown in Table 2.

**Table 2.** Performance of 7 dimension reduction methods on training dataset PDB1075.

| Algorithm | ACC (%) | SN (%) | SP (%) | MCC |
| --- | --- | --- | --- | --- |
| LLE | 78.26 | 82.82 | 73.95 | 0.5690 |
| SVD | 82.10 | 83.59 | 80.69 | 0.6426 |
| SVM-RFE | 90.82 | 88.22 | 93.26 | 0.8167 |
| LASSO | 91.75 | 89.38 | 93.99 | 0.8354 |
| Elastic net | 92.60 | 91.12 | 93.99 | 0.8519 |
| LinearSVC | 92.03 | 91.70 | 92.35 | 0.8405 |
| XGB-RFE | 93.44 | 93.44 | 93.44 | 0.8687 |

It can be seen from Table 2 that XGB_RFE has the best performance among the 7 dimension reduction methods. The values of ACC and MCC both reach the highest which are 93.44% and 0.8687 respectively. The ACC value of XGB_RFE is 15.18%, 11.34%, 2.62%, 1.69%, 0.84% and 1.41% higher than LLE, SVD, SVM-RFE, LASSO, Elastic net and LinearSVC respectively. The MCC value of XGB_RFE is 29.97%, 22.61%, 5.20%, 3.33%, 1.68% and 2.82% higher than LLE, SVD, SVM-RFE, LASSO, Elastic net and LinearSVC respectively. ROC and PR curves can more intuitively compare the performance of 7 different feature selection methods in Supplementary Figure S2. From the above analysis, it shows that XGB-RFE can reduce model complexity while eliminating redundant and irrelevant features. It can also improve model accuracy and shorten model running time. Therefore, we choose XGB-RFE as the dimension reduction method and finally get the 100-dimension optimal feature.

### 3.4. Feature visualization

The distribution of the Fusion feature and the optimal feature (Fusion (XGB-RFE)) are shown in the feature space to explain that XGB-RFE can improve prediction accuracy. For comparison, the original feature space and the optimal feature space are converted to a two-dimension space by T-distributed Stochastic Neighbor Embedding (t-SNE) [78]. The t-SNE visualization is shown in Fig. 3.

**Fig. 3.**
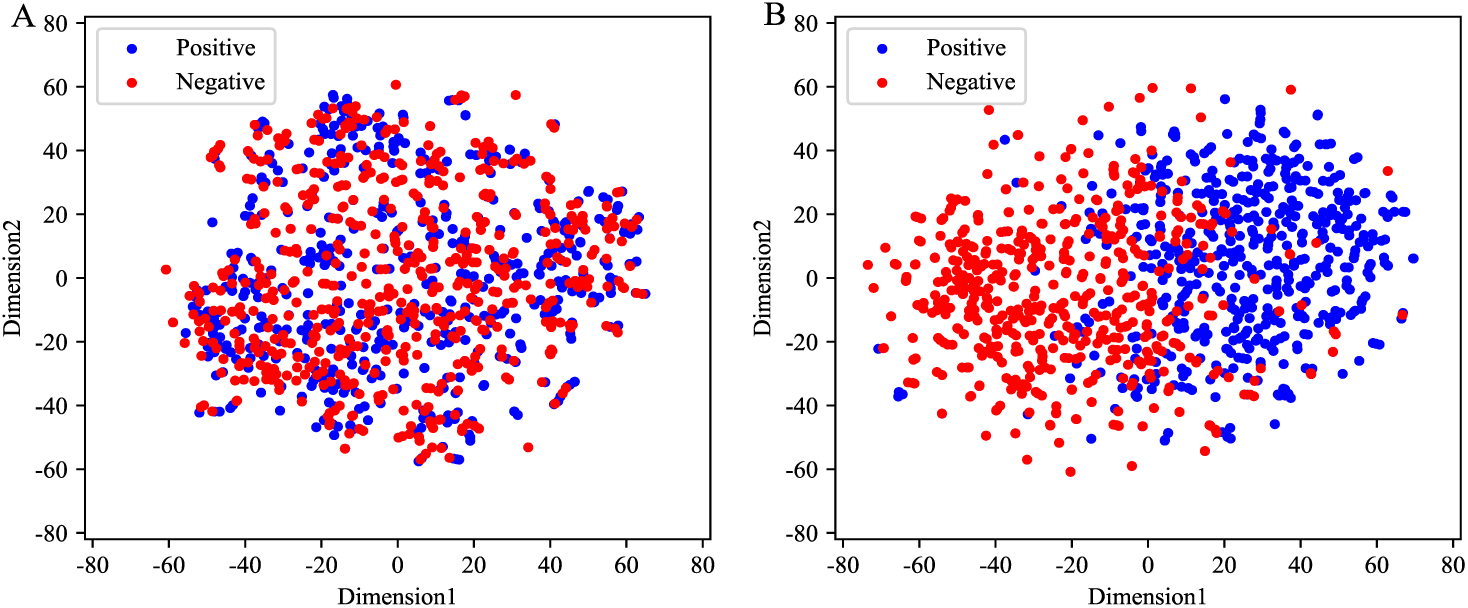
The t-SNE visualization of the Fusion feature (A) and Fusion (XGB-RFE) features (B) in two-dimension space.

It can be seen from Fig. 3 (A) that the positive and negative examples of the Fusion feature are mixed in a two-dimension space. There is no obvious distinction between the positive examples and negative examples, which brings greater challenges to the prediction of DBPs. Compared with the distribution of Fusion features, the distribution of positive and negative samples in Fusion (XGB-RFE) features is more obvious from Fig. 3 (B). The positive and negative examples are gathered in different areas in the two-dimension space, which can capture the difference between the positive and negative samples. Also, XGB-RFE is effective in transforming features from high-dimension space to low-dimension space, which can shorten training time. It can provide more effective information for the identification of DBPs and improve the prediction accuracy of the model.

### 3.5. Selection of base-classifier

To determine the most suitable classifier, 9 machine learning classifiers are tested. The parameters of 9 machine learning classifiers are as follows, i.e., (1) The closest neighbor of KNN is 5; (2) SVM uses the RBF kernel function; (3) RF sets the number of base decision trees to 500 and the maximum learning depth to 10; (4) The number of GBDT iterations is 500; (5) The number of iterations of XGBoost is 500; (6) AdaBoost sets the number of base decision trees to 500; (7) The number of iterations of LightGBM is 500; and (8) NB and LR use default parameters. The prediction results of 9 classifiers on the training dataset PDB1075 are as Table 3.

**Table 3.** Performance of 9 base-classifiers on the training dataset PDB1075.

| Model | ACC (%) | SN (%) | SP (%) | MCC |
| --- | --- | --- | --- | --- |
| NB | 65.60 | 36.68 | 92.90 | 0.3600 |
| KNN | 75.54 | 66.02 | 84.52 | 0.5156 |
| RF | 83.51 | 84.92 | 82.15 | 0.6707 |
| LR | 83.88 | 81.47 | 86.16 | 0.6775 |
| SVM | 84.72 | 85.33 | 84.15 | 0.6945 |
| AdaBoost | 86.41 | 84.56 | 88.16 | 0.7280 |
| GBDT | 86.69 | 84.17 | 89.07 | 0.7339 |
| XGBoost | 90.07 | 88.42 | 91.62 | 0.8013 |
| LightGBM | 92.59 | 89.59 | 95.45 | 0.8528 |

In Table 3, the ACC of NB, KNN, RF, LR, SVM, AdaBoost, GBDT, XGBoost and LightGBM are 65.60%, 75.54%, 83.51%, 83.88%, 84.72%, 86.41%, 86.69%, 90.07%, and 92.59%, respectively. The ACC of LightGBM is 26.99% and 17.05% higher than that of NB and KNN. The ACC values of LightGBM and XGBoost classifiers both exceed 90%. XGBoost is only 2.52% lower than LightGBM. The MCC of LightGBM and XGBoost are 0.8528 and 0.8013, respectively. LightGBM is 0.4928 higher than NB on MCC, and XGBoost is 0.4413 higher than NB on MCC.

The ROC and PR curves can more vividly represent the performance of 9 different classifiers, as shown in Fig. 4. In Fig. 4, the AUC of LightGBM is 0.9758, which is the highest among 9 base-classifiers. The area covered by ROC curve of XGBoost is second-largest with an AUC value of 0.9638. From Fig. 4 (B), the AUPR value of LightGBM is largest which is 0.9781. The AUPR value of XGBoost is second-largest which is 0.9663. Considering the performance of 9 base-classifiers, XGBoost and LightGBM have high accuracy and stability. Thus, XGBoost and LightGBM are selected as the best combination of base-classifier.

**Fig. 4.**
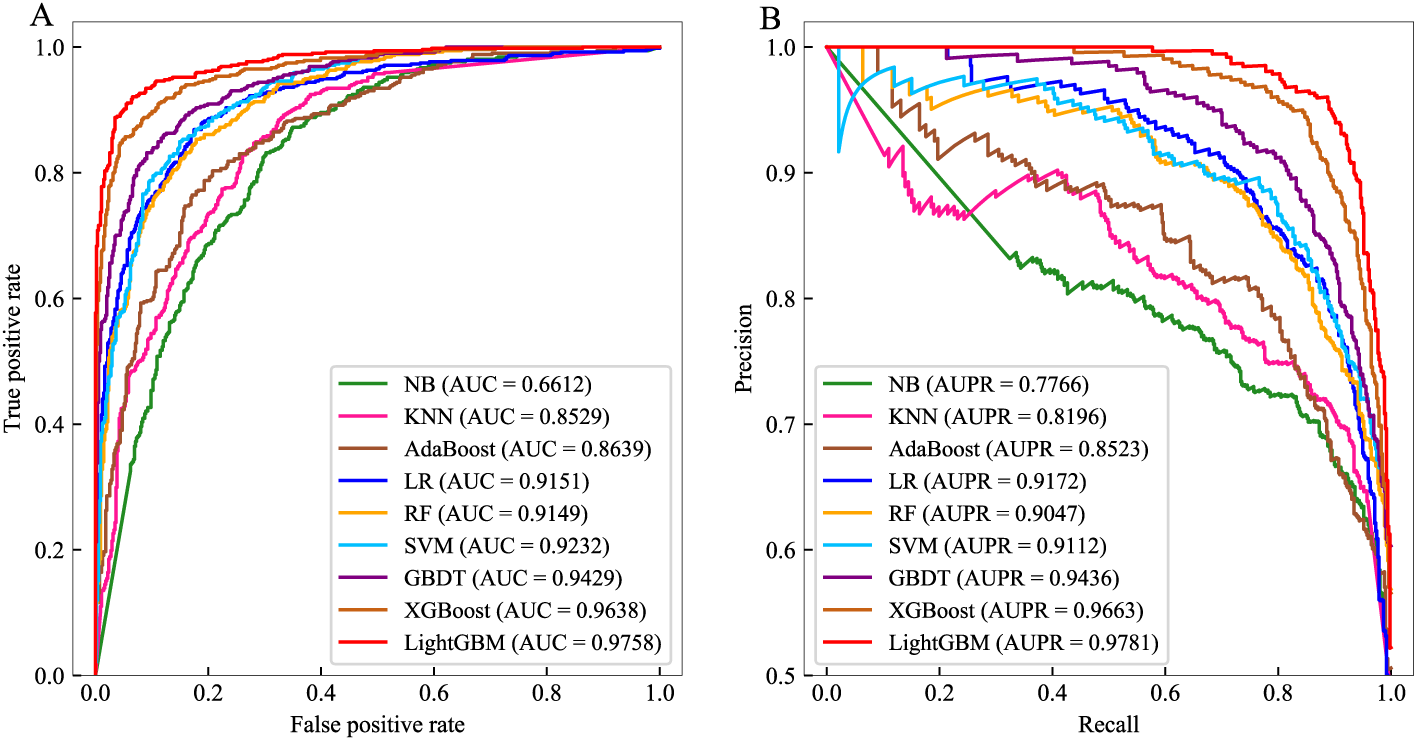
ROC and PR curve of different base-classifiers on the training dataset PDB1075.

### 3.6. Selection of meta-classifier

After the training on the first stage, the binding probability and non-binding probability of each protein sequence are obtained from LightGBM and XGBoost. The output probability is input into the meta-classifier for training again. Therefore, the choice of meta-classifier also plays a significant role in the model establishment. The specific parameters of 9 classifiers are as follows, (1) the number of XGBoost iterations is 500; (2) The base-classifier of AdaBoost and GBDT both select decision trees (500); (3) LightGBM iterates 500 times; (4) The number of KNN neighbors is 5; (5) SVM uses the RBF kernel function; (6) The base decision trees number of RF is 500 and the maximum learning depth as 10; and (7) NB and LR use default parameters. The performance of 9 meta-classifiers is shown in Table 4.

**Table 4.** The performance of 9 meta-classifiers on the training dataset PDB1075.

| Model | ACC (%) | SN (%) | SP (%) | MCC |
| --- | --- | --- | --- | --- |
| NB | 89.50 | 89.58 | 89.44 | 0.7900 |
| XGBoost | 88.38 | 91.70 | 85.25 | 0.7698 |
| AdaBoost | 89.03 | 93.05 | 85.25 | 0.7838 |
| LightGBM | 88.75 | 91.12 | 86.52 | 0.7763 |
| KNN | 89.03 | 92.66 | 85.61 | 0.7833 |
| LR | 89.41 | 88.42 | 90.35 | 0.7880 |
| GBDT | 89.60 | 93.24 | 86.16 | 0.7946 |
| RF | 90.07 | 93.44 | 86.89 | 0.8036 |
| SVM | 93.44 | 93.44 | 93.44 | 0.8687 |

In Table 4, SVM outperforms 9 classifiers. SVM has 93.44% ACC, which is 4.41%, 4.03%, 3.84%, and 3.37% higher than KNN, LR, GBDT, and RF respectively. The MCC of SVM is 0.8687, which is 7.87%, 9.89%, 8.49% and 9.24% higher than NB, XGBoost, AdaBoost and LightGBM respectively. The combination of SVM, XGBoost, and LightGBM increases the diversity of the stacked ensemble classifier and obtains better prediction results. We further evaluate the performance of the 9 meta-classifiers through ROC and PR curves, as shown in Fig. 5.

**Fig. 5.**
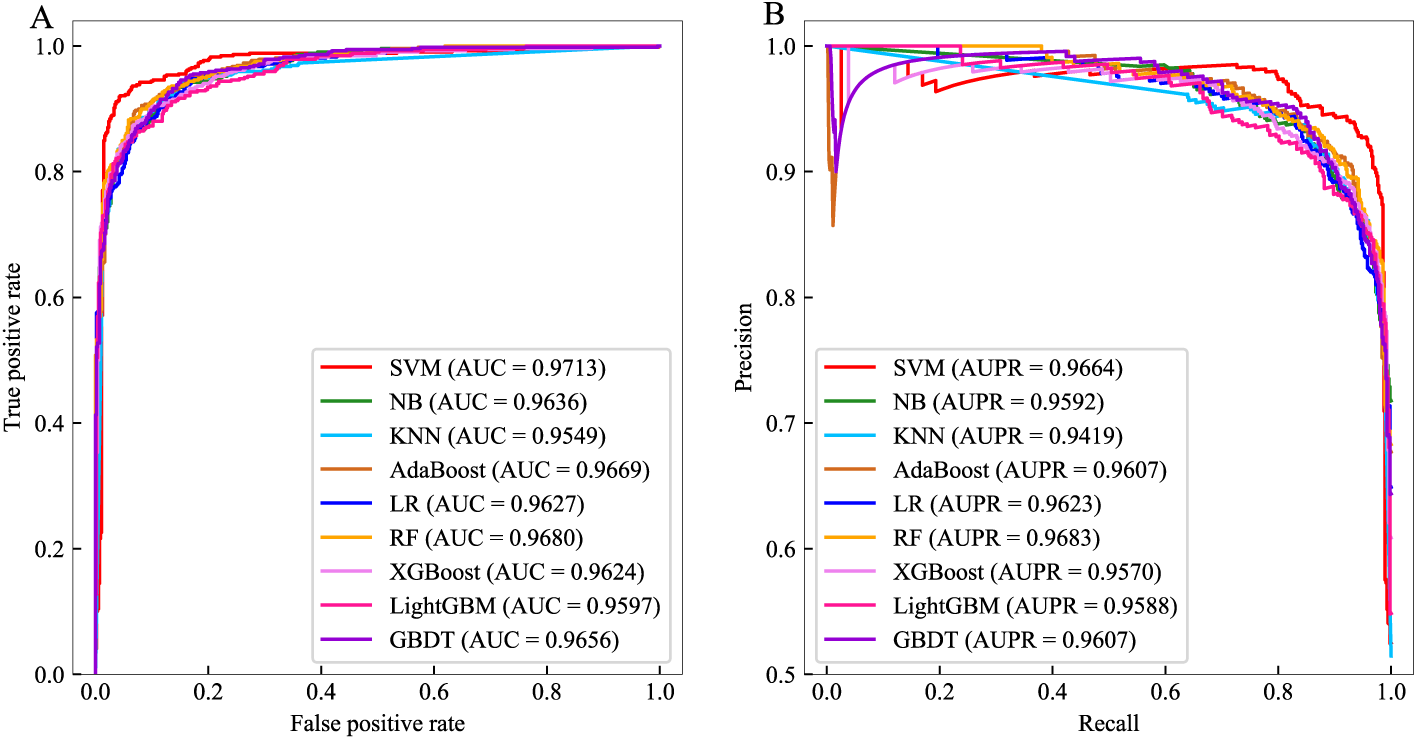
The ROC and PR curves of 9 meta-classifiers on the training dataset PDB1075.

In Fig. 5., the area covered by ROC curve of SVM is maximal with an AUC value of 0.9713. The AUC value of SVM is 0.33%-1.64% higher than NB, KNN, AdaBoost, LR, RF, XGBoost, LightGBM, and GBDT (0.9731 vs. 0.9636, 0.9549, 0.9669, 0.9627, 0.9680, 0.9624, 0.9597, 0.9656). The area covered under the PR curve of the SVM is 0.9664, which is 0.0019 lower than the AUPR value of RF. The AUPR value of SVM is 0.41%-2.45% higher than NB, KNN, AdaBoost, LR, RF, XGBoost, LightGBM, and GBDT (0.9664 vs. 0.9592, 0.9419, 0.9607, 0.9623,0.96 83, 0.9570, 0.9588, 0.9607). Comparing with other classifiers, SVM shows strong predictive ability. SVM realizes the mapping from low-dimension space to high-dimension space by RBF function. The optimal hyperplane is found in the high-dimension space to distinguish between DBPs and non-DBPs. Thus, SVM is selected as a meta-classifier.

### 3.7. Comparison with other state-of-the-art methods

To verify the effectiveness of StackPDB, StackPDB is compared with PSSM-DT [33], HMMBinder [79], iDNAPro-PseAAC [80], DBPPred-PDSD [17], iDNAProt-ES [11], HMMPred [13], Local-DPP [28], DP-BINDER [26]. PSSM-DT [33] proposed a new feature extraction method PSSM distance transformation (PSSM-DT) and combined with SVM to predict DBPs. HMMBinder [79] used monogram features and bigram features for feature extraction which converted HMM matrix into the same length vectors. Then the feature vectors were input into SVM to construct the HMMBinder model. iDNAPro-PseAAC [80] extracted protein sequence features based on physicochemical properties and evolutionary information and used SVM to construct iDNAPro-PseAAC. Table 5 shows the comparison of StackPDB and other published methods.

**Table 5.** Comparison of StackPDB with other DBPs prediction methods on the training set PDB1075 based on the LOOCV.

| Methods | ACC (%) | SN (%) | SP (%) | MCC |
| --- | --- | --- | --- | --- |
| iDNAPro-PseAAC [80] | 76.56 | 75.62 | 77.45 | 0.5300 |
| Local-DPP [28] | 79.20 | 84.00 | 74.50 | 0.5900 |
| PSSM-DT [33] | 79.96 | 81.91 | 78.00 | 0.6220 |
| HMMPred [13] | 83.90 | 83.98 | 83.82 | 0.6800 |
| HMMBinder [79] | 86.33 | 87.07 | 85.55 | 0.7200 |
| DBPPred-PDSD [17] | 89.02 | 89.14 | 88.88 | 0.7800 |
| iDNAProt-ES [11] | 90.18 | 90.38 | 90.00 | 0.8000 |
| DP-BINDER [26] | 92.46 | 91.80 | 93.07 | 0.8400 |
| StackPDB | **93.44** | 93.44 | 93.44 | **0.8687** |

In Table 5, the ACC of StackPDB reaches 93.44%, which is 16.88%, 14.24%, 13.48%, 9.54%, 7.11%, 4.42%, 3.26% and 0.98% higher than the ACC values of iDNAPro-PseAAC, Local-DPP, PSSM-DT, HMMPred, HMMBinder, DBPPred-PDSD, iDNAProt-ES and DP-BINDER, respectively. The MCC of StackPDB is 0.8687, which exceeds the MCC values of iDNAPro-PseAAC, Local-DPP, PSSM-DT and HMMPred by 33.87%, 27.87%, 24.67%, and 18.87% respectively. The histogram of StackPDB compared with other DBPs prediction methods is shown in Supplementary Figure S3. Compared with other 8 published methods, StackPDB performs the best.

To evaluate the predictive ability of StackPDB more fairly and objectively, PDB186 and PDB180 are applied to verify our StackPDB. Then the test results are compared with several published methods. The feature extraction parameters, dimension reduction method, and classifier parameters of the independent test datasets are consistent with the training set, which can make the test results more rigorous and reliable. Considering the validity of the comparison results, the test results of the independent test set PDB186 are compared with those already published methods HMMPred [13], HMMBinder [79], DBPPred [35], Local-DPP [28], PSSM-DT [33], MSFBinder [30] and iDNAProt-ES [11]. Compared the test results of the independent test set PDB180 with competitive DNAbinder [27], DNA-Prot [81], iDNA-Prot [82] and Top-2-gram-SVM [36]. DBPPred [35] extracted features based on sequence information, solvent accessibility, secondary structural information, and evolutionary information. RF was used to feature selection. Finally, Gaussian Navïe Bayes (GNB) was used to predict DBPs. Top-2-gram-SVM [36] combined PseAAC and top-n-grams to extract evolutionary information and physicochemical properties. Finally, the classifier SVM was used to predict DBPs. DNA-Prot [81] extracted the physicochemical properties and secondary structural information of protein sequences and used RF to predict DBPs. iDNA-Prot [82] was proposed by Lin et al., using grey system theory to improve PseAAC and choosing RF for DBPs prediction. The comparison results are shown in Table 6 and Table 7.

**Table 6.** Comparison of the independent test dataset PDB186 with other state-of-art methods under the verification of the LOOCV method.

| Methods | ACC (%) | SN (%) | SP (%) | MCC |
| --- | --- | --- | --- | --- |
| HMMBinder [79] | 69.02 | 61.53 | 76.34 | 0.3900 |
| DBPPred [35] | 76.90 | 79.60 | 74.20 | 0.5380 |
| Local-DPP [28] | 79.00 | 92.50 | 65.60 | 0.6250 |
| PSSM-DT [33] | 80.00 | 87.09 | 72.83 | 0.6470 |
| MSFBinder [30] | 80.11 | 92.47 | 67.74 | 0.6200 |
| HMMPred [13] | 81.18 | 94.62 | 67.74 | 0.6480 |
| iDNAProt-ES [11] | 80.64 | 81.31 | 80.00 | 0.6100 |
| StackPDB | **84.40** | 83.87 | 84.95 | **0.6882** |

**Table 7.** Comparison of the independent test dataset PDB180 with other state-of-art methods under the verification of the LOOCV method.

| Methods | ACC (%) | SN (%) | SP (%) | MCC |
| --- | --- | --- | --- | --- |
| DNAbinder [27] | 78.89 | 54.32 | 98.98 | 0.6100 |
| DNA-Prot [81] | 76.67 | 66.67 | 84.85 | 0.5300 |
| iDNA-Prot [82] | 81.11 | 72.84 | 87.88 | 0.6200 |
| Top-2-gram-SVM [36] | 85.56 | 82.72 | 87.88 | 0.7100 |
| StackPDB | **90.00** | 91.36 | 88.89 | **0.7997** |

In Table 6, the ACC value of StackPDB on PDB186 exceeds other prediction methods. The ACC of StackPDB is 84.40%, which is 3.22%-15.38% higher than the ACC of HMMBinder, DBPPred, Local-DPP, PSSM-DT, MSFBinder, HMMPred, and iDNAProt-ES (84.40 vs. 69.02, 76.90, 79.00, 80.00, 80.11, 81.18, 80.64). From the perspective of model stability, the MCC of StackPDB is 0.6882, which is 29.82%-4.02% higher than the MCC of HMMBinder, DBPPred, Local-DPP, PSSM-DT, MSFBinder, HMMPred, and iDNAProt-ES (0.6882 vs. 0.39, 0.5380, 0.6250, 0.647, 0.62, 0.648, 0.61). It can be seen that the StackPDB model also has high stability. As we can see from Table 7, the prediction results of the StackPDB are better than other methods.

The ACC value of the StackPDB model reached 90.00%, which is 11.11%, 13.33%, 8.89% and 4.44% higher than DNAbinder, DNA-Prot, iDNA-Prot and Top-2-gram-SVM respectively. The MCC value reaches 0.7997, which is 18.97%, 26.97%, 17.97% and 8.97% higher than DNAbinder, DNA-Prot, iDNA-Prot and Top-2-gram-SVM respectively. Supplementary Figure S4 and Figure S5 shows the histograms of the independent test datasets PDB186 and PDB180 compared with other DBPs prediction methods. The performance of StackPDB on the independent test datasets PDB186 and PDB180 show that the StackPDB model not only has the high predictive ability but also shows great potential in the generalization ability and stability. Hence, StackPDB is a competitive predictor of DBPs.

## 4. Conclusion

DBPs not only play a significant role in human life activities but also guide the development of disease treatment and drug research and development. With the rapid growth of DBPs, the development of DBPs prediction models has become a central issue in bioinformatics. We propose a new method, called StackPDB. First, five feature extraction methods extract the information, where PsePSSM, EDT, RPT, and PSSM-TPC extract evolutionary information. Especially, PSSM-TPC extracts the evolutionary information. PseAAC can effectively obtain the physicochemical properties information. Fusion of five features can obtain different aspects of protein sequence information. Second, we use XGB-RFE to decrease the feature dimension. XGB-RFE combines the gradient boosting and recursive feature elimination, which can fully learn the importance score of each feature. It can also eliminate redundant and irrelevant features without losing important features and reduce the complexity of the model. The final predictor of DBPs is stacked ensemble classifier which composed of XGBoost, LightGBM and SVM. Stacked ensemble classifier can take advantage of multiple classifiers, reduce generalization errors, and have stronger predictive ability than ordinary machine learning classifiers. StackPDB has achieved good prediction results on the training dataset PDB1075 based on LOOCV. Compared with other state-of-art methods, StackPDB shows strong predictive ability on the independent test set PDB186 and PDB180. In future work, deep learning methods are considered to predict DNA binding proteins. Deep learning has powerful fitting capabilities and can approximate any complex function. In particular, it has a great advantage in processing data with a large sample size, which can make better accuracy of DBPs prediction.

## Declaration of competing interest

No author associated with this paper has disclosed any potential or pertinent conflicts which may be perceived to have impending conflict with this work.

## Acknowledgments

This work was supported by the National Nature Science Foundation of China (No. 61863010), the Key Research and Development Program of Shandong Province of China (No. 2019GGX101001), and the Natural Science Foundation of Shandong Province of China (No. ZR2018MC007, ZR2019MEE066).

## Notes

### Competing Interest Statement

The authors have declared no competing interest.

