## Supplementary Tables, Supplementary Figures for "StackPDB: predicting DNA-binding proteins based on XGB-RFE feature optimization and stacked ensemble classifier"

**Table of contents**

**1. Supplementary Method Illustration**

**Si1.** K nearest neighbor (KNN) classifier

**Si2.** Support vector machine (SVM)

**Si3.** Random forest (RF) classifier

**Si4.** Naïve bayes (NB) classifier

**Si5.** Logistic regression (LR)

**Si6.** Adaptive boosting (AdaBoost)

**Si7.** Gradient boosting decision tree (GBDT)

**Si8.** LightGBM

**Si9.** Extreme gradient boosting (XGBoost)

**2. Supplementary Tables**

**Table S1.** Performance of PseAAC with different parameters on the training dataset PDB1075.

**Table S2.** Performance of PsePSSM with different parameters on the training dataset PDB1075.

**3. Supplementary Figures**

**Figure S1.** ROC and PR curves of different feature extraction methods over LOOCV.

**Figure S2.** ROC and PR curve of different dimension reduction methods.

**Figure S3.** Comparison of training dataset PDB1075 with other DBPs predicting methods over LOOCV.

**Figure S4.** Comparison of independent test dataset PDB186 with other DBPs predicting methods over LOOCV.

**Figure S5.** Comparison of independent test dataset PDB180 with other DBPs predicting methods over LOOCV.

### 4. Supplementary References

### 1. Supplementary Method Illustration

To find the base-classifier used in the first stage and the meta-classifier used in the second stage, we explore 9 different machine learning algorithms.

#### Si1. K nearest neighbor (KNN) classifier

We use KNN [1] as one of the candidate methods of base-classifier and meta-classifier. Given a training dataset $T=\{(x_1,y_1),(x_2,y_2),\cdots,(x_N,y_N)\}$, KNN finds $k$ neighbors closest to the instance in the training dataset for a new input instance $x$. According to the voting principle, the majority of these $k$ instances belong to a certain class $y_i$ and then $x$ is divided into class $y_i$.

#### Si2. Support vector machine (SVM)

We use SVM [2] as one of the candidate methods of base-classifier and meta-classifier. Given a training dataset $T=\{(x_1,y_1),(x_2,y_2),\cdots,(x_N,y_N)\}$, the learning strategy of SVM is to find an optimal hyperplane based on the training dataset and separate different samples. Assuming that the hyperplane is $\omega^T x+b=0$, the distance from any point $x$ to the hyperplane $(\omega,b)$ in the sample space is $r=\frac{|\omega^T x+b|}{\|\omega\|}$. To maximize the interval, it can be transformed into the solution of formula (1).

$$\begin{aligned}&\min_{\omega,b}\frac{1}{2}\|\omega\|^2\\&s.t.\ \ y_i(\omega^T x_i+b)\geq 1,\ \ i=1,2,\cdots,m\end{aligned}\tag{1}$$

#### Si3. Random forest (RF) classifier

We use RF [3] as one of the candidate methods of base-classifier and meta-classifier. Based on Bagging integration, RF further adds random attribute selection in the training process of the decision tree. For each node of the base decision tree, a subset containing $k$ attributes is randomly selected from the attribute set of the node, and then an optimal attribute is selected from this subset for the division.

#### Si4. Naïve bayes (NB) classifier

We use NB [4] as one of the candidate methods of base-classifier and meta-classifier. NB is a classification method based on Bayes' theorem and the independence assumption of characteristic conditions. First, for a given training dataset $T=\{(x_1,y_1),(x_2,y_2),\cdots,(x_N,y_N)\}$, the joint probability distribution of input or output is learned based on the assumption of independent feature conditions.

For a given input $x$, the output $y$ with the largest posterior probability is found according to Bayes' theorem.

### Si5. Logistic regression (LR)

We use LR [5] as one of the candidate methods of base-classifier and meta-classifier. LR uses logistic regression to generate an estimated probability to measure the relationship between the dependent variable (in this study: whether the protein is DBPs or not) and one or more independent variables. Grid search is used to optimize the parameter $C$ that controls the regular strength to achieve the highest accuracy based on LOOCV.

### Si6. Adaptive boosting (AdaBoost)

We use AdaBoost [6] as one of the candidate methods of base-classifier and meta-classifier. AdaBoost is a typical Boosting algorithm, which is based on the "additive model" that is the linear combination of the base learner to minimize the exponential loss function [7].

$$\ell_{\exp}(H \mid D) = E_{x\sim D}\left[e^{-f(x)H(x)}\right] \tag{2}$$

The AdaBoost model is as follows:

$$H(x) = sign\left(\sum_{t=1}^{T} \alpha_t h_t(x)\right) \tag{3}$$

### Si7. Gradient boosting decision tree (GBDT)

We use GBDT [8] as one of the candidate methods of base-classifier and meta-classifier. GBDT can also be expressed as $H(x) = \sum_{t=1}^{T} \alpha_t h_t(x)$ based on the "additive model". Through continuous iteration, the true value of the sample is fitted to the residual of the current classifier to approximate the true value. The prediction result of the $m$-th base-classifier is:

$$F_t(x) = F_{t-1}(x) + \alpha_t h_t(x) \tag{4}$$

The optimization goal of $h_t(x)$ is to minimize the gap between the current prediction results $F_{t\text{-}1}(x_i) + h(x_i)$ and $y_i$.

$$h_t = \arg\min_h \sum_{i=1}^{n} L(y_i, F_{m-1}(x_i) + h(x_i)) \tag{5}$$

### Si8. LightGBM

We use LightGBM [9] as one of the candidate methods of base-classifier and meta-classifier. LightGBM is an efficient gradient boosting decision tree. Its base-classifier is a decision tree that can be trained sequentially by fitting the negative gradient of the loss function. It uses gradient-based unilateral sampling (GOSS) and proprietary feature bundling (EFB). GOSS is used to divide the optimal node by calculating variance gain, while EFB reduces the number of effective features by bundling mutually exclusive features to speed up the training process.

**Si9. Extreme gradient boosting (XGBoost)**

We use XGBoost [10] as one of the candidate methods of base-classifier and meta-classifier. XGBoost is an ensemble algorithm based on gradient boosting. XGBoost is an optimization model that combines a linear model with a boosting tree model. It uses not only the first derivative of the loss function but also the second derivative of the loss function. Given a training dataset $T=\{(x_1,y_1),(x_2,y_2),\cdots,(x_N,y_N)\}$, the objective function of the XGBoost algorithm is:

$$obj(\theta)=\sum_{i}^{n} l(y_i,\hat{y}_i)+\sum_{t=1}^{T}\Omega(f_t) \tag{6}$$

According to the Taylor expansion optimization of the objective function, the second-order Taylor expansion of the loss function after $t$ iterations is obtained, and the approximate loss function is obtained:

$$L^{(t)}=\sum_{i=1}^{k}\left[l(y_i,\hat{y}^{(t-1)})+g_i f_t(x_i)+\frac{1}{2}h_i f_t^2(x_i)\right]+\Omega(f_t) \tag{7}$$

### 2. Supplementary Tables

**Table S1.**

Performance of PseAAC with different parameters on the training dataset PDB1075.

| $\lambda$ | 5 | 10 | 15 | 20 | 25 | 30 | 35 | 40 | 45 |
|---|---|---|---|---|---|---|---|---|---|
| ACC (%) | 73.95 | 75.07 | 73.95 | 73.76 | 73.76 | 73.95 | 72.91 | 72.07 | 71.51 |
| SN (%) | 74.71 | 74.71 | 72.20 | 71.24 | 74.90 | 75.68 | 75.68 | 71.04 | 72.97 |
| SP (%) | 73.22 | 75.41 | 75.59 | 76.14 | 72.68 | 72.31 | 70.31 | 73.04 | 70.13 |
| MCC | 0.4791 | 0.5011 | 0.4783 | 0.4745 | 0.4756 | 0.4798 | 0.4600 | 0.4409 | 0.4309 |

**Table S2.**

Performance of PsePSSM with different parameters on the training dataset PDB1075.

| $\xi$ | 1 | 2 | 3 | 4 | 5 |
|---|---|---|---|---|---|
| ACC (%) | 76.66 | 77.41 | 77.23 | 76.76 | 76.76 |
| SN (%) | 83.01 | 81.85 | 80.89 | 80.69 | 81.47 |
| SP (%) | 70.67 | 73.22 | 73.77 | 73.04 | 72.31 |
| MCC | 0.5398 | 0.5519 | 0.5472 | 0.5381 | 0.5391 |
| $\xi$ | 6 | 7 | 8 | 9 | 10 |
| ACC (%) | 76.85 | 76.76 | 77.04 | 76.94 | 77.13 |
| SN (%) | 82.05 | 81.27 | 82.24 | 80.69 | 80.50 |
| SP (%) | 71.95 | 72.50 | 72.13 | 73.41 | 73.95 |
| MCC | 0.5417 | 0.5389 | 0.5455 | 0.5417 | 0.5450 |

### 3. Supplementary Figures

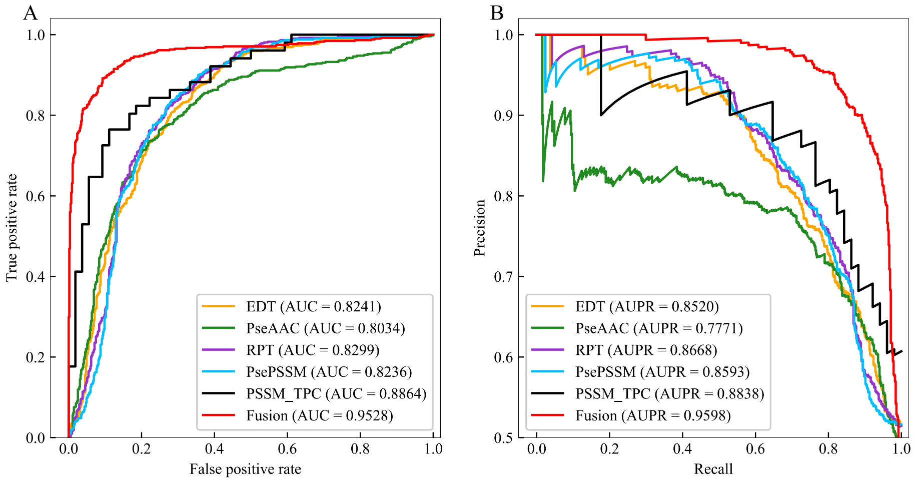


**Figure S1.** ROC and PR curves of different feature extraction methods over LOOCV.

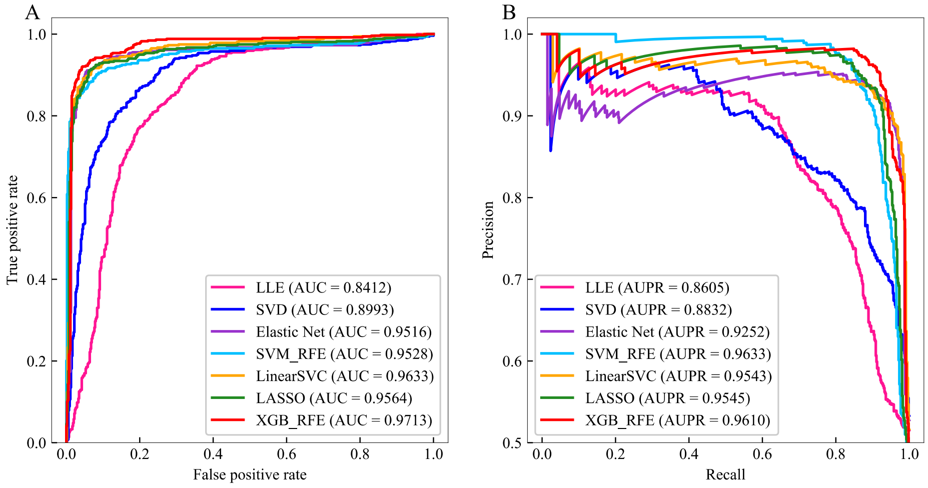


**Figure S2.** ROC and PR curve of different dimension reduction methods.

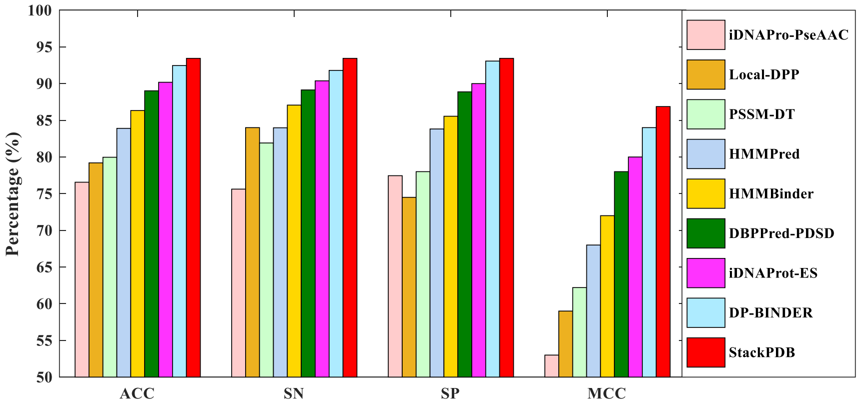

**Figure S3.** Comparison of training dataset PDB1075 with other DBPs predicting methods over LOOCV.

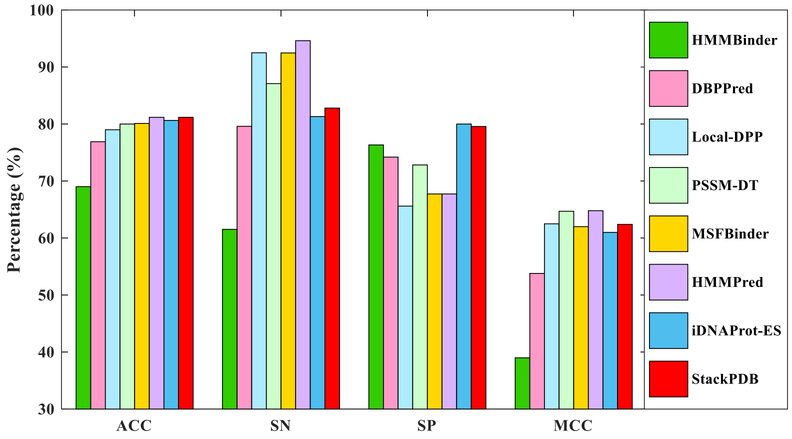


**Figure S4.** Comparison of independent test dataset PDB186 with other DBPs predicting methods over LOOCV.

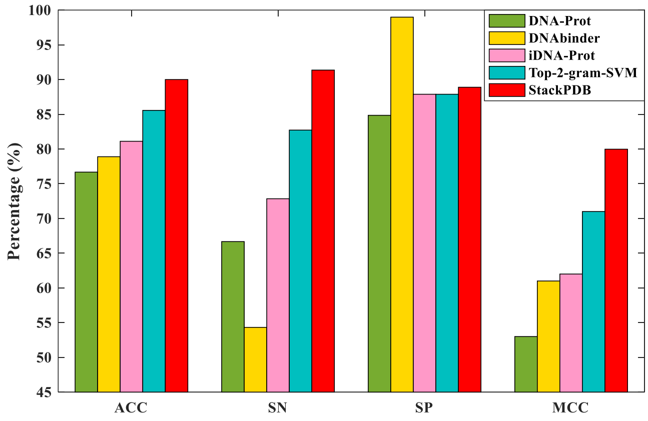


**Figure S5.** Comparison of independent test dataset PDB180 with other DBPs predicting methods over LOOCV.
